# Quantifying growth perturbations over the fattening period in swine via mathematical modelling

**DOI:** 10.1101/2020.10.22.349985

**Authors:** Revilla Manuel, Lenoir Guillaume, Flatres-Grall Loïc, Muñoz-Tamayo Rafael, Nicolas C Friggens

## Abstract

**Background:** Resilience can be defined as the capacity of animals to cope with short-term perturbations in their environment and return rapidly to their pre-challenge status. In a perspective of precision livestock farming, it is key to create informative indicators for general resilience and therefore incorporate this concept in breeding goals. In the modern swine breeding industry, new technologies such as automatic feeding system are increasingly common and can be used to capture useful data to monitor animal phenotypes such as feed efficiency. This automatic and longitudinal data collection integrated with mathematical modelling has a great potential to determine accurate resilience indicators, for example by measuring the deviation from expected production levels over a period of time.

**Results:** This work aimed at developing a modelling approach for facilitating the quantification of pig resilience during the fattening period, from approximately 34 kg to 105 kg of body weight. A total of 13 093 pigs, belonging to three different genetic lines were monitored (Pietrain, Pietrain NN and Duroc) since 2015, and body weight measures registered (approximately 11.1 million of weightings) with automatic feeding systems. We used the Gompertz model and linear interpolation on body weight data to quantify individual deviations from expected production, thereby creating a resilience index (*ABC*). The estimated heritabilities of *ABC* are low but not zero from 0.03 to 0.04 (± 0.01) depending on the breed.

**Conclusions:** Our model-based approach can be useful to quantify pig responses to perturbations using exclusively the growth curves and should contribute to the genetic improvement of resilience of fattening pigs by providing a resilience index.

## Background

Climate change and societal concerns (*e.g*., animal welfare and use of antibiotics) on livestock production result in important challenges for animal breeding. Alternatives to address these challenges include the implementation of strategies to select animals that can adapt to a changing environment and to promote a healthy environment for facilitating farm management (1). In this context, the last decade has seen an enormous increase in interest in animal robustness to environmental effects. Friggens *et al*. define the robustness as the ability, in the face of environmental constraints, to carry on doing the various things that the animal needs to do to favour its future ability to reproduce (2). Concomitantly, the concept of resilience has emerged in animal sciences encompassing not only the response of the individual to diseases challenge but also the individual’s response to other sources of stressors. Colditz and Hine defined resilience as the capacity of the animal to be minimally affected by disturbances or to rapidly return to the state pertained before exposure to a disturbance (3). Several definitions and resilience-associated concepts have been discussed in literature (1), reflecting the interest of this concept in a broad range of scientific disciplines (4).

In the era of big data collection on farms, the digitalization process will generate new knowledge in most of the relevant topics in swine production including nutrition, health management, reproduction, genetics, biosecurity, behavior, welfare, and pollutant emissions (5). Sensors (6), such as commercially available automatic feeding systems (AFS), capture longitudinal data (feed intake -FI-, feeding time, daily visits and body weight -BW-). These data can be further exploited using the knowledge of animal requirements and physiology to develop new phenotypes increasing sustainability and efficiency of breeding. Such an exploitation calls for adequate mathematical tools. AFS allow pigs to feed *ad libitum* and recognize individual growing pigs via a radio frequency identification (RFID) transponder. The large number of automatic BW registers measured by AFS could generate knowledge for management decision-making. In particular, the detection of BW deviations from standard trajectories would generate useful insights on the status of animal with minimum effort if automated.

Animal breeding is showing an increasing interest for resilience to be included as a trait in breeding goals. However, the incorporation of resilience in swine breeding goals is currently an uncommon practice. One of the main drawbacks that hinder the incorporation of resilience in breeding is the difficulty of providing quantitative resilience indicators (2). Recent technological developments based on longitudinal data give new opportunities to define resilience indicators based on the difference between observed production and an individual’s potential production (although the definition of the individual potential is a challenging issue). Several studies have explored continuous recording of pig performance to study the impact of perturbations, including novel phenotypes related to disease resilience using daily FI (7, 8), and modelling approaches to detect potential perturbations as deviations of FI (9). Modelling efforts to characterize the animal response to perturbations in dairy cattle have also been developed (10). Our group has recently developed a modelling approach, for facilitating the quantification of piglet resilience to weaning (11). In our previous work, we proposed a resilience indicator that has the potential to be used in elite breeding populations. Building upon our previous work, the aim of the present study was to develop a modelling methodology for quantifying an individual pig resilience indicator based on longitudinal BW measurements registered routinely by an AFS during the fattening period. Moreover, the genetics underlying this resilience indicator were analyzed in two of the most used commercial breeds to show the potential to improve resilience of swine livestock through inclusion of this indicator in breeding goals.

## Methods

### Data source

The pigs used in this study belonged to the Piétrain (Pie) and Duroc (Du) pure breeds. Piétrain is an European sire line breed, strongly selected for lean meat content during the last decades (12). The Du breed is also used as a terminal sire when fattening pigs are produced. The Du breed has both an excellent growth rate and high intramuscular fat (13). AXIOM Genetics have two different lines belonging to Piétrain breed namely Piétrain Français NN Axiom line (Pie NN) with pigs free from halothane-sensitivity and Piétrain Français Axiom line with animals positive to this gene.

A total of 13 093 boars belonging to three different lines were used in this study: 5 841 and 5 032 belonging to Pie and Pie NN line respectively, and 2 220 belonging to Du breed.

### Station conditions

The boar testing station of the breeding company AXIOM Genetics (Azay-sur-Indre, France), built in 2015, located in the Centre region in France housed the animals used in this study. A group of 336 piglets were introduced to the station every 3 weeks. AXIOM’S requirements for biosafety are applied: forward march, showers and change of clothes, cleaning and disinfection program, blood monitoring. The boars arrived, after weaning, from 7 different birth farms (5 farms for Pie, 1 farm for Pie NN and 1 farm for Du) to the herd when they were between 25 and 35 days of age (8 ± 3 kg BW). Birth farms are integrated into the AXIOM breeding scheme, comply with AXIOM’s biosafety and health requirements (monitoring, vaccination plan) and are negative for major diseases. For each batch, all pigs arrived within 1 successive week and were kept in the same pen of 14 animals. Each pen is made up of 14 male piglets from the same breed and from the same birth farm. The composition of the pens is never modified, with no reallocation. They were kept in air-filtered quarantine rooms (nursery) for 5 weeks, the time needed for seroconversion control and to validate there are not positive to major disease, such as porcine reproductive and respiratory syndrome (PRRS), brucellosis, swine influenza, etc. They were then raised in post-weaning rooms for 2 weeks. The three lines are present in each group in the station and meet the same breeding conditions. Then they were transferred in fattening rooms when they were approximately between 70 and 80 days of age (34.4 kg). They were kept in fattening rooms for 65 to 77 days until the individual testing (weighing, ultrasonic backfat and muscle measurements) around 150 days of age (104 kg BW). Animals were kept in the same pen from arrival until slaughter. The station consisted in 2 nursery rooms, 2 post-weaning rooms and 10 fattening rooms with 12 identical pens each, housing a maximum of 14 pigs per pen, leading to a total capacity of 2 638 pig places. Only fattening rooms are equipped with AFS. Each pen had one water nipple available for the animals. One group, from the same week of introduction in the station, is divided in two fattening rooms (24 pens with 14 pigs).

### Automatic individual body weight data collection

An AFS pig performance testing feeding station (Nedap N.V.; Groenlo, the Netherlands) was located in the front of each of the pen. The feeder was 0.7 m wide, and the total length was 1.69 m. The feeder included a feed trough and had no gates. The feeder only allows the entrance of one animal. The pig entering the feeder was individually identified via an electronic RFID transponder located in the ear. All animals were maintained under standard intensive rearing conditions and were fed individually ad libitum from the feeder with a standard diet non limiting in amino-acids. Briefly, the growing diet provided 9.75 MJ/kg of net energy with 15% of crude protein and 0.9% of lysine. The boars were not castrated.

Data collection started when animals were transferred in fattening pens and finished 1 week after individual testing. Animals were individually weighted the day of transfer (IW: initial weight) and the day of individual testing (WT).

The data analyzed in this study used information registered at each visit in the AFS on individual pigs relating to identification number, date, location, duration of the visit, FI and BW. The dataset included boars raised at the station from September 2015 to July 2019. During that period, 65 batches arrived at the station (13 093 pigs in total).

### Data pre-treatment

Datasets were processed separately for the three lines. Each dataset from the AFS was thoroughly assessed in order to validate the data, and identify important data gaps and quality issues using SAS (14). The different datasets were analyzed independently but using the same procedure.

In the raw data file, one record corresponded to one animal visit to an AFS. A first processing step consisted of eliminating the records without an RFID tag detected, and without a valid association between animal ID and RFID tag.

As a second step of quality control for each visit, the weight was considered as null for records without BW record, with a duration of the feeder visit <5s (scale stabilization) and for weights measured during the 6 first days of the fattening period that were out of a range between 0.7*IW and 1.3*IW. Indeed, during the first 6 days, the pigs are in the adaptation phase and the AFS stalls remain open. It is possible that two pigs try to enter in the AFS stall at the same time or that a pig puts a leg in the feeder causing an incorrect weight measurement.

For the third control step, a quadratic regression of weight on age + age^2^ for each animal was applied to eliminate aberrant weights. The ratio between the residual value and the fitted value was calculated for each visit of each animal. If the ratio was > 0.15, the measured weight was considered to be null. The ratio of 0.15 was selected by using a trial-and-error approach to find a compromise between the data cleaning and the number of data points to be kept for further analysis. This step was repeated a second time excluding the initially identified aberrant weights. Following this step, the visits of an animal during a day were aggregated in a single record. The weight of the day was estimated from the median of the non-null weights (WM) measured during the day’s visits. If the number of non-null weights for the day was <3, the median of daily weights was considered to be null.

The fourth control step consisted in analyzing all of data from each AFS within fattening group (AFS*Group) in order to detect inconsistencies linked to the AFS machine. A linear regression of WM on days (number of days since the beginning of measurements) was applied. The standard deviation of the residual value was calculated for each day for each AFS*Group. If more than 10% of the weights measured on AFS*Group were > 3 * standard deviation, then AFS*Group records have been removed from the data set. The objective was to rule out animals from AFS with a mechanical problem. Animals with less than 15 AFS measurements in total or more than 10 consecutive days without measurements were removed from the analysis. We accepted that animals had missing weights during the fattening period.

The total FI (TFI) during control period was calculated as the sum of FI for all visits during the control period. When a control day is missing (i.e., due to a mechanic problem of AFS or loss of a RFID tag), the missing daily FI is estimated by using local regression, “*proc loess*” implement in SAS (14).

Finally, for visualization purpose a kernel density estimation was performed to produce a smoothed color representation of a scatterplot by using the “*smooothScatter*” function implement in R (15). Multivariate kernel smoothing is described by Wand and Jones (16).

### Two-step mathematical model approach

Our modelling approach comprises two steps. The first step looks at determining a theoretical (potential) growth curve of each animal. The second step looks at constructing the actual perturbed growth curve. The resulting two curves are the ingredients for further determination of an individual resilience indicator.

Animal growth models aim at describing the pattern of growth over the animal’s lifetime, defining an upper limit to growth. In our study, we assumed that, under ideal conditions, animal growth follows the theoretical (potential) growth of the animal not experiencing any perturbation. The potential growth of each pig was modelled using the Gompertz equation (i) (17), using the formulation described on Schulin-Zeuthen *et al*. (18).

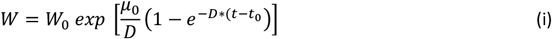

where *W*_0_ is the value of live weight *W* (kg) at the initial time of the recordings (*t*_0_), *μ*_0_ (d^-1^) is the initial value of the specific growth rate at *t*_0_, the constant *D* (d^-1^) is a growth rate coefficient that controls the slope of the growth rate (*μ*) curve and *t* (days) is time since birth. All parameters are positive. In the remaining text, we will call the trajectories that resulted from this calibration as the unperturbed curve. The unperturbed growth model resulted in two parameters to be estimated, *μ_0_* and *D*. As explained below in the model calibration section, we constructed the unperturbed curve such that the perturbed data cannot be above the unperturbed curve by a margin of 5%. The value of 5% was set in accordance with the accuracy provided by AFS.

For our second modelling step, since the Gompertz equation is a monotonic function that does not account for possible decrease of BW due to perturbations, we construct a perturbed growth curve using the daily BW measurements registered routinely by the AFS. For missing records, values were estimated using the linear interpolation method implemented in the “*interp1*” function in Scilab (19). It should be noted that if high frequency data are available, the linear interpolation step is not needed.

We further calculated the difference of the area under the curve between the perturbed curve and the unperturbed growth. The area under the curve was calculated using the trapezoidal rule implemented in the “*inttrap*” Scilab function. The resulting value was called Area Between Curves (*ABC*) index, and was considered as a proxy of resilience (the lower *ABC* the higher the resilience or an animal faced to low perturbation). For those non-normal distributed values, the *ABC* parameter results were normalised applying the log_2_ transformation. Visualization of the quartiles distribution of this parameter was performed with the ‘*ggridges*’ R package (15).

Finally, correlation analyses were performed to explore the relationships between growth model parameters to be estimated (*μ*_0_, *D*) and *ABC*. Pearson correlations were analyzed in R using the ‘*cor*’ function in the base package.

### Model calibration

The parameters *μ*_0_, *D* of the Gompertz model for each animal were estimated by minimizing the normalized least square error with a penalized function (ii):

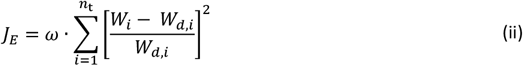

where *W_d_* is the weight data (kg), *W* the weight predicted by the model, and *n_t_* the total number of measurements. The parameter *ω* is a penalization factor that we constructed to constrain the unperturbed curve to envelope all experimental data. The penalization factor is defined as follows (iii):

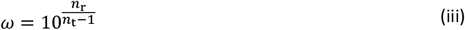

Where *n_t_* is the number of measurements for each animal and *n*_r_ is the number of records where the ratio between the residual (real BW – predicted weight) and the real BW was higher than 5%.

### Phenotypic swine production traits

When the average weight of the group was approximately 100 kg, the individual testing was performed. Measurements made during the test were: weight (WT), average ultrasonic backfat thickness (BF: mean of 3 measurements) and ultrasonic *longissimus dorsi* thickness (LD: 1 measurement). BF and LD were adjusted to 100 kg live weight (BF100 and LD100 respectively) by applying linear coefficients. These equations are based on those established by Jourdain *et al*. (20).

The average daily gain (ADG), expressed in g/day, was calculated as the ratio between the BW gain (WG), difference between WT and IW, and number of days of control period. The feed conversion ratio (FCR) was calculated as the ratio between TFI during the fattening period and WG, expressed in kg/kg.

The selection traits estimated in the 3 lines are BF100, LD100, ADG and FCR.

### Statistical analyses

For each breed, the ABC, BF100, LD100, ADG and FCR traits were first analyzed separately with a linear mixed model (LMM). The global statistical model was defined as (iv):

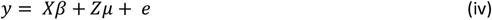

where *y* is the vector of phenotype measures for a trait, *β* is the vector of fixed effects depending on the trait considered (Table S1). *X* is the known matrix for fixed effects. *μ* is the vector of animal genetic random effects with ~ N(0, A σ^2^u) where A is the pedigree-based relationship matrix. Z is the known design matrices for animal genetic effect. *e* is a vector of residual random effects with e ~ N(0, I σ2e) where **I** is the identity matrix of appropriate size.

Variance components (variance and covariance) were estimated using the REML method with ASReml 3.0 (21) separately for each line.

Heritability was calculated as the ratio of animal genetic variance to the phenotypic variance. Due to convergence issues, correlations between *ABC* and selection traits were estimated using two-trait analyses for lines Pie and Pie NN. Genetic correlations have not been estimated for Duroc due to insufficient data.

For Pie, 24 generations of pedigree information comprising 57 459 animals from 1991 to 2019 were included in the analysis. For Pie NN, 24 generations of pedigree information comprising 16 137 animals from 1993 to 2019 were included in the analysis. For Du, 22 generations of pedigree information comprising 20 632 animals from 1995 to 2019 were included in the analysis.

## Results

### Data pre-treatment procedure

From a total of 13 093 animals, more than 11.1 million measurements (1 measurement = 1 visit including BW and FI recording) were registered using the AFS. These numbers correspond to the raw dataset. We implemented a data pre-treatment procedure to provide high quality data for the modelling approach. This dataset was analyzed separately in three different data subsets belonging to Pie, Pie NN and Du breed lines, and the same procedure was applied in each dataset. The comparison between the number of animals in the filtered data and the raw dataset showed a ratio of 0.93, 0.91 and 0.87, for Pie, Pie NN and Du lines, respectively. Regarding the number of AFS measurements, the ratios between the filtered and the raw dataset were 0.76 for Pie, 0.69 for Pie NN, and 0.77 for Du. Complete descriptive statistics for the dataset used in this study are shown in Table 1

**Table 1.**
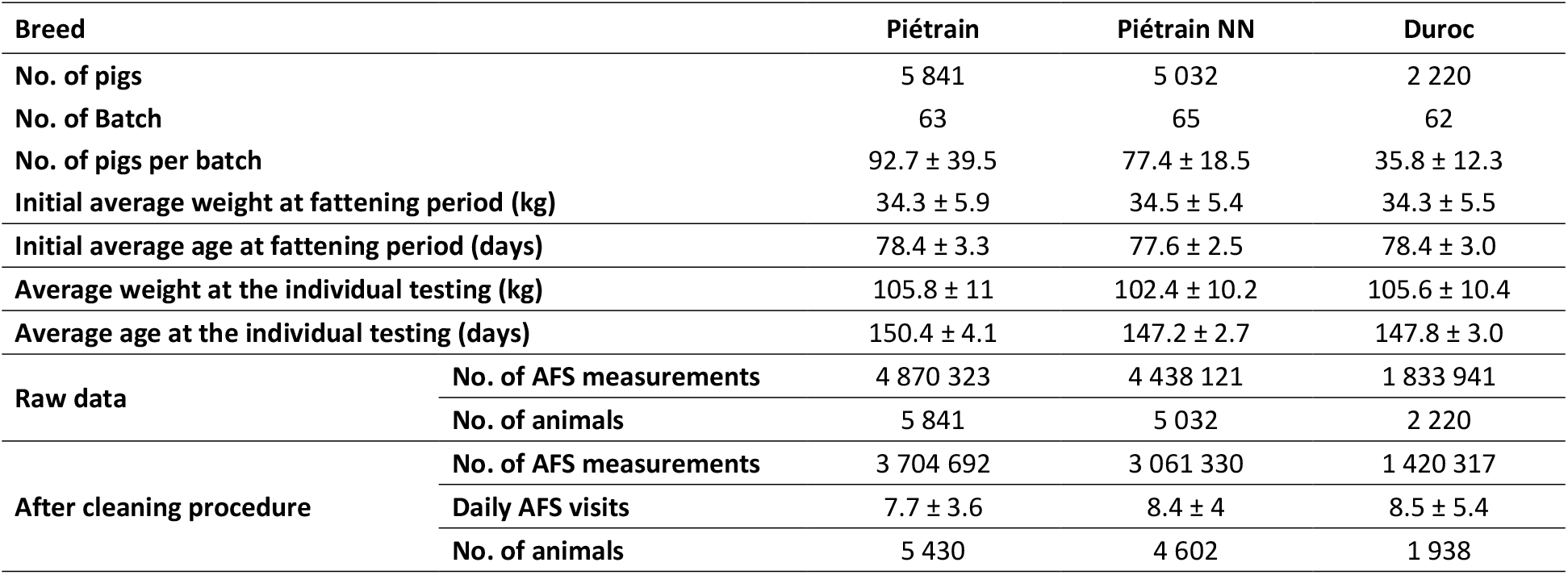
Descriptive statistics for the datasets used in this study.

The data analyzed in this study included information from a total of 11 970 boars, belonging to three of the most common lines used in swine industry. The final data set consisted of daily median BW records from 409 770, 337 964, and 140 170 Pie, Pie NN and Du measurements, respectively.

A visual comparison of the AFS measurements dataset of Pie line before and after the data cleaning procedure is shown in Figure 1. Moreover, a graphic representation of Pie NN and Du lines filtering procedure is shown in Figure S1 and S2, respectively. The figure illustrates the proportion of measurement points discarded from the analysis before filtering (Figure 1: A1-A4; Figure S1: A1-A4 and Figure S2: A1-A4 - Raw data) and after filtering (Figure 1: B1-B4; Figure S1: B1-B4 and Figure S2: B1-B4 - Filtered data), especially weights with a value close to zero.

**Figure 1.**
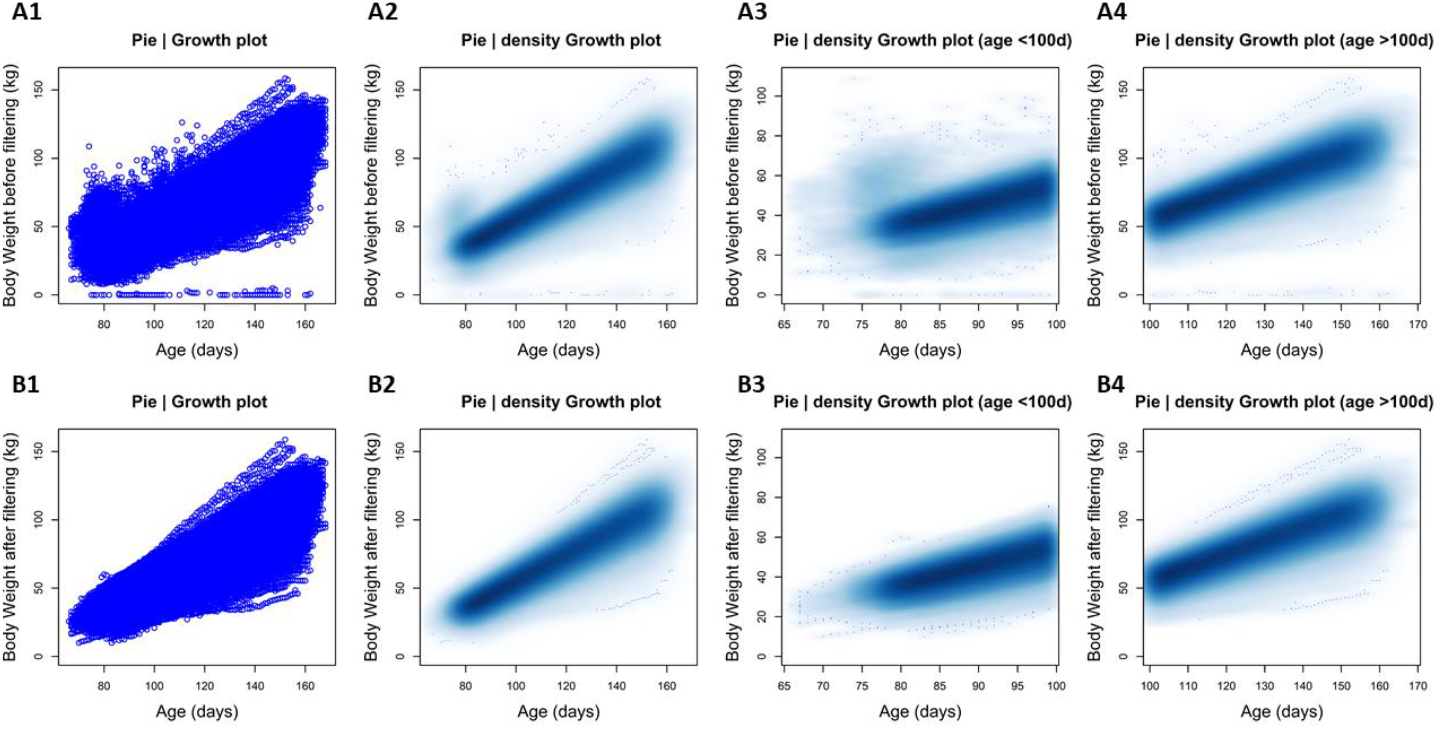
Comparison of body weight density plots before (A) and after (B) applying data cleaning procedure in Pie line. In **A1** and **B1** plots each point represent the median of the individual daily body weight registered by the AFS during the pig fattening period. **A2** and **B2** are smoothed color density representations of a scatterplot. Shaded areas are constructed to illustrate the density of points falling into each part of the plot allowing for an intuitive visualization of very large datasets. A zoom in the density scatter plot before (**A3-B3**) and after (**A4-B4**) 100 days of individual age is illustrated.

### Growth curve modelling over pig fattening period

To quantify the deviation of the unperturbed curve from the perturbed curve, we constructed the parameter *ABC* as a resilience indicator. Figure 2 displays the BW dynamic trajectories of two animals belonging to Pie line exhibiting different patterns. For an animal with a growth performance close to the unperturbed model (Figure 2A), *ABC* was 37 657. For an animal with high degree of perturbation (Figure 2B), *ABC* was 493 007. The parameter ABC is a useful indicator of the degree of perturbation of an animal and allows comparison within a population. Table 2 summarizes the complete descriptive statistics of the model parameters.

**Figure 2.**
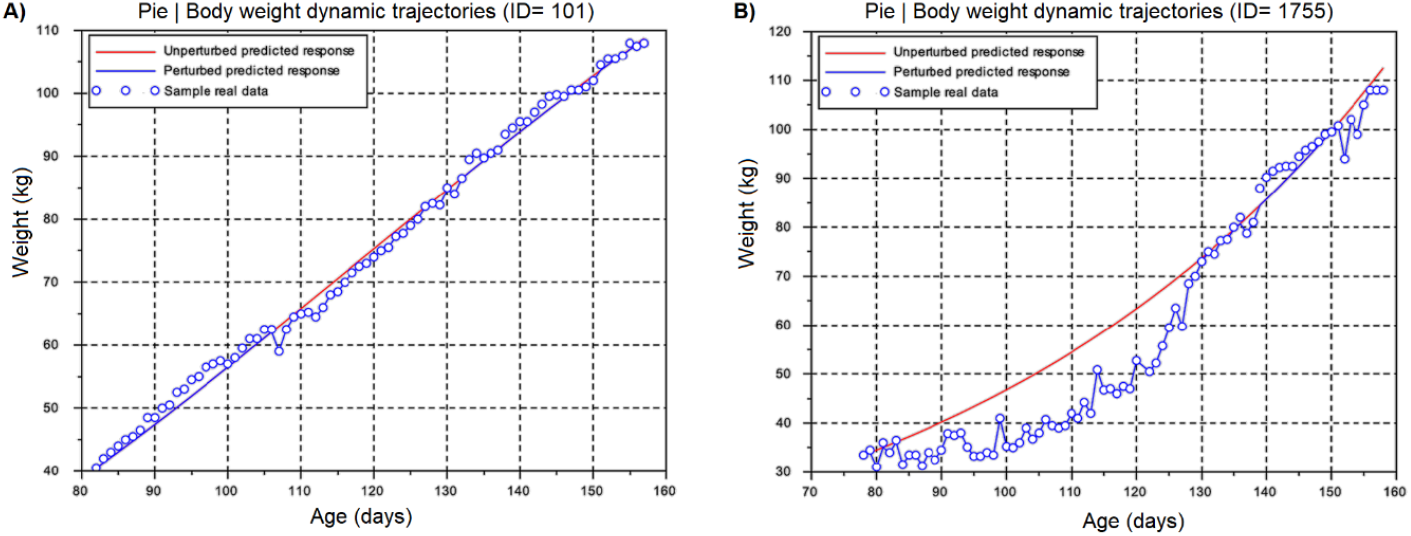
Comparison of the perturbed (blue line) and the unperturbed (red line) predicted response based on the body weight dynamic trajectories recorded during the whole fattening period. Circles represent the median daily body weight measures of the individual pig. Two different animals belonging to Pie line are represented.

**Table 2.** Descriptive statistics of the parameters for the growth curve modelling in the three pig lines analyzed.

| Breed |  | Piétrain | Piétrain NN | Duroc |
| --- | --- | --- | --- | --- |
| $\mu_0$ | Range | $7.13 \times 10^{-04} - 1.00 \times 10^{-01}$ | $2.79 \times 10^{-03} - 9.34 \times 10^{-02}$ | $6.58 \times 10^{-03} - 8.46 \times 10^{-02}$ |
| | Mean | $2.62 \times 10^{-02}$ | $2.59 \times 10^{-02}$ | $2.78 \times 10^{-02}$ |
| | SD | $1.95 \times 10^{-02}$ | $6.19 \times 10^{-03}$ | $8.40 \times 10^{-03}$ |
| $D$ | Range | $1.18 \times 10^{-16} - 7.19 \times 10^{-01}$ | $1.00 \times 10^{-09} - 2.66 \times 10^{-01}$ | $1.03 \times 10^{-15} - 6.52 \times 10^{-02}$ |
| | Mean | $1.16 \times 10^{-02}$ | $1.64 \times 10^{-02}$ | $1.65 \times 10^{-02}$ |
| | SD | $2.22 \times 10^{-02}$ | $8.83 \times 10^{-03}$ | $7.85 \times 10^{-03}$ |
| ABC parameter | Min. | 239 | 2 253 | 2 788 |
|  | 1<sup>st</sup> quartile | 26 244 | 27 489 | 31 518 |
|  | Median | 33 564 | 35 804 | 44 069 |
|  | Mean | 41 556 | 46 474 | 58 441 |
|  | 3<sup>rd</sup> quartile | 44 524 | 49 257 | 69 738 |
|  | Max | 703 283 | 595 914 | 407 425 |

$\mu_0$ : individual growth rate ( $d^{-1}$ ); $D$ : extent of the exponential decay of the growth ( $d^{-1}$ ); ABC: area between the perturbed and the unperturbed growth curves; SD: standard deviation.

Furthermore, Figure 3 represents a visual comparison of the model parameters for the three analyzed lines. Parameter *μ_0_* (Figure 3A) showed no significant differences when Pie and Pie NN lines were analyzed, nevertheless both of them were significantly different (*p*-value ≤ 0.001) compared with Du line. In the case of parameter *D* (Figure 3B) significant differences were found between Pie and Pie NN (*p*-value ≤ 0.001), and Pie and Du lines (*p*-value ≤ 0.05).

**Figure 3.**
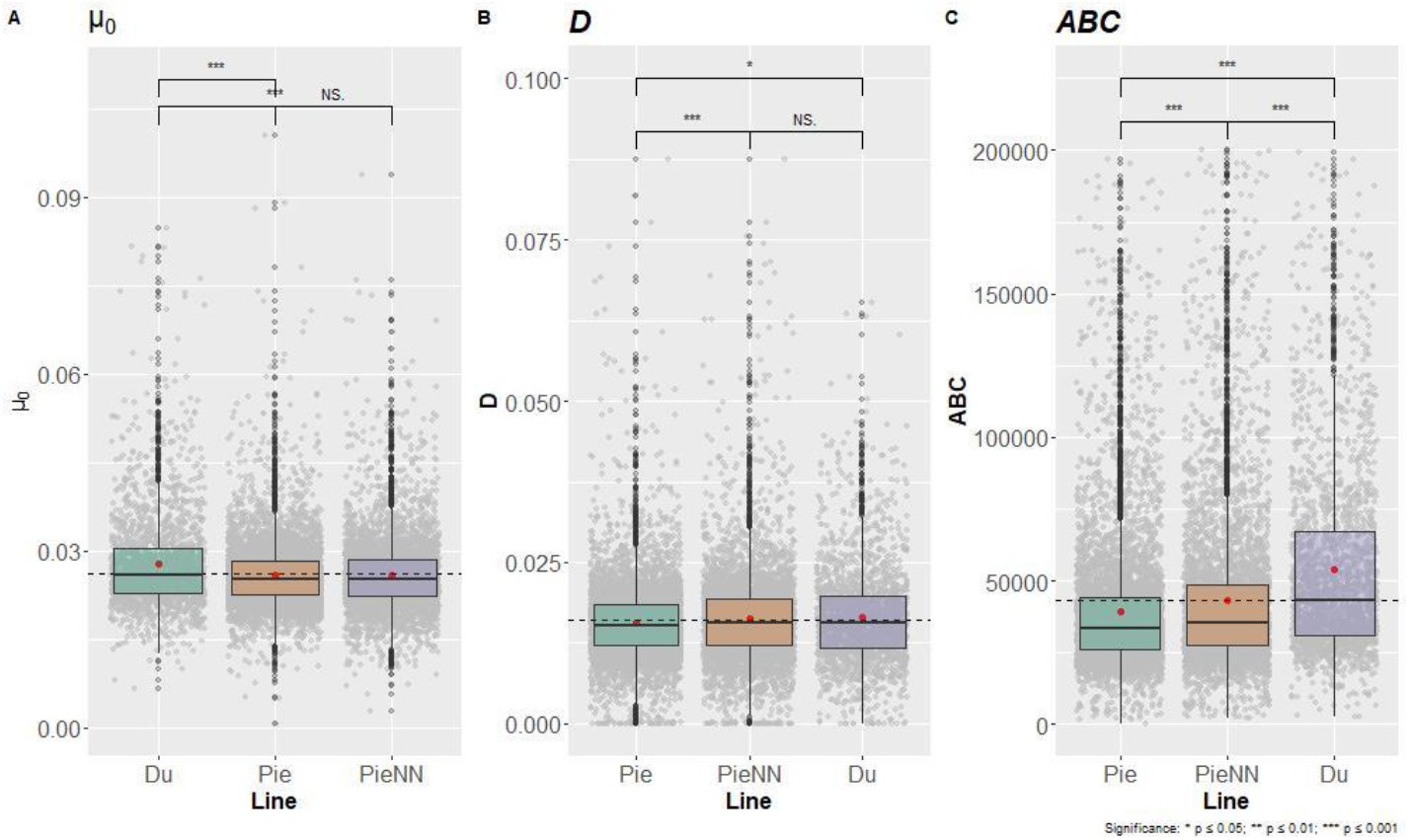
Comparison of *μ_0_, D and ABC* statistics in the three pig lines analyzed. Parameter *μ_0_* (**A - initial growth rate value**), parameter *D* (**B – exponential rate of decay of growth rate**), and parameter *ABC* (**C – area between the perturbed and unperturbed growth curves**) are represented Red points show the average value of the model parameters for each line. The dotted line represents the global average of the parameter. Significant differences between groups are indicated as \**p*-value≤ 0.05, \*\**p*-value≤ 0.01, and \*\*\**p*-value≤ 0.001.

For the parameter *ABC* (Figure 3C) significant differences were identified in all the comparisons performed (*p*-value ≤ 0.001). Despite the observed significant differences for the parameter *ABC*, their distribution between Pie and Pie NN lines were similar (Figure 4A and 4B), compared with the distribution observed for Du line (Figure 4C). This result is logical due to the close genetic origin of both lines.

**Figure 4.**
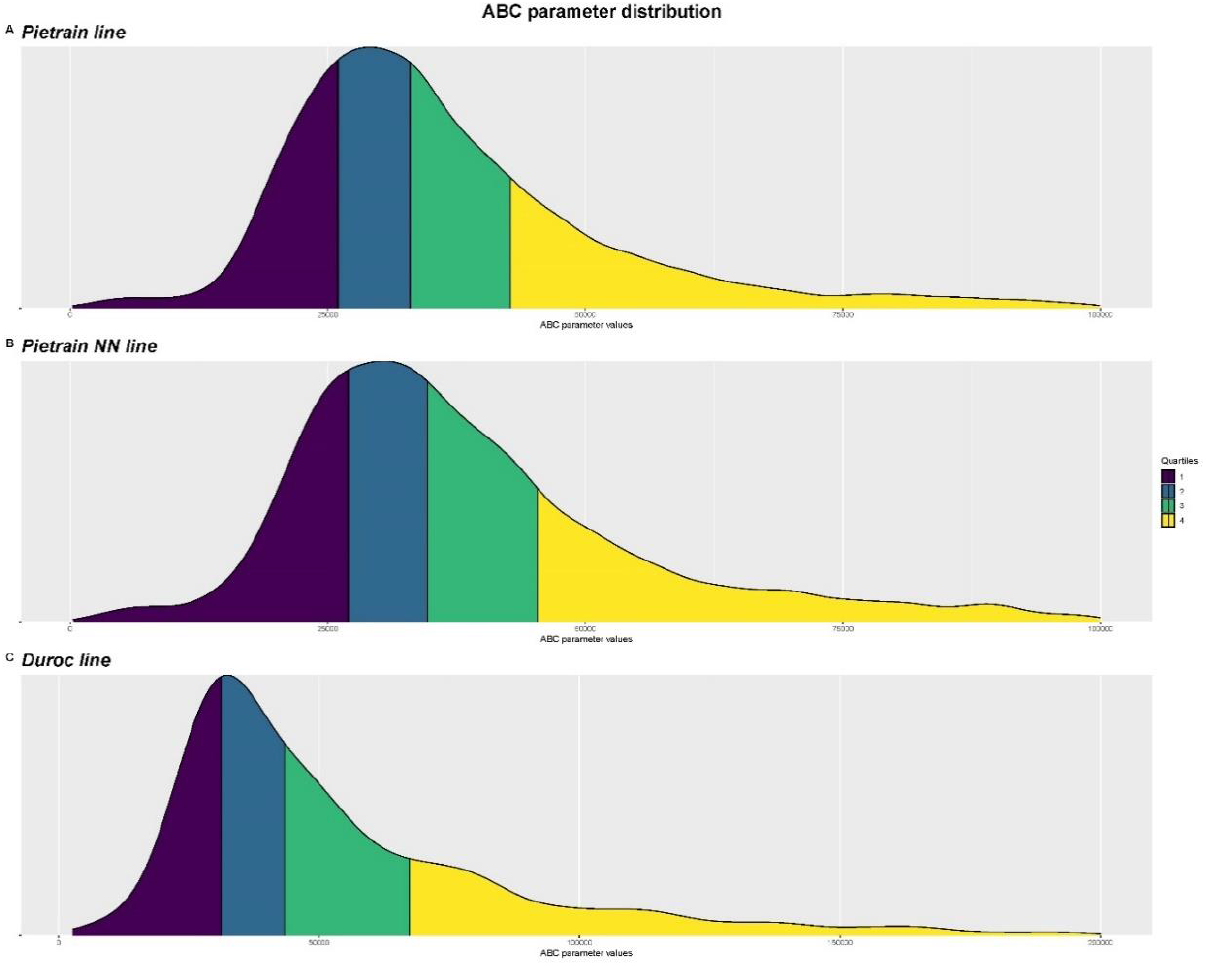
*ABC* parameter distribution in the three pig lines analyzed. *ABC*: area between the perturbed and the unperturbed growth curves. Colors represent quartiles information.

Moreover, correlations between the model parameters of the three lines were analyzed (Table 3). The parameter *μ_0_* showed positive significant correlations with parameter *D* in the three analyzed lines, 0.88 for Pie, 0.81 for Du, and 0.62 for Pie NN. In the case of parameter *μ_0_* and parameter *ABC* significant correlations were only identified in Du (0.37) and Pie NN lines (0.20). A similar pattern was also identified between parameter *D* and parameter *ABC*, being Du (0.30) and Pie NN (0.19) lines those that showed significant correlations.

**Table 3.**
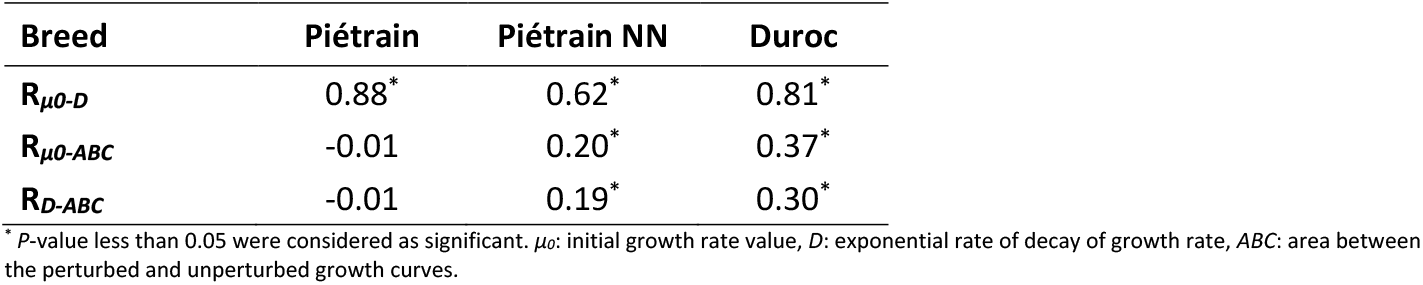
Pearson’s correlation coefficients among the growth curve model parameters in the three pig lines analyzed.

### Estimating trait heritability and genetic correlations

The heritability of the *ABC* parameter was analyzed (Table 4), ranging between 0.03 and 0.04. Both pig breeds had similar heritability. Phenotypic and genetic correlations were also performed between the *ABC* parameter and important swine production traits such as BF100, LD100, ADG and FCR (Table 5). Phenotypic correlations between *ABC* and production traits are close to 0 for both breeds, ranging from −0.09 to 0.10. Genetic correlations between *ABC* and production traits are low to moderate. In both breeds, the highest genetic correlation is between the resilience index and ADG, with values of 0.59 for Pie and 0.39 for Pie NN.

**Table 4.** Estimated heritabilities (*h*^2^) and corresponding standard errors (SE) of *ABC* parameter in Pie and Pie NN.

| Breed | Piétrain | Piétrain NN |
| --- | --- | --- |
| $h^2$ (SE) | 0.04 ± 0.01 | 0.03 ± 0.016 |

**Table 5.** Estimates of heritabilities (diagonal) and of genetic (above diagonal) and phenotypic (below diagonal) correlations among *ABC* and four commercial selection traits in Pie and Pie NN.

| Breed | Piétrain |  |  |  |  | Piétrain NN |  |  |  |  |
| --- | --- | --- | --- | --- | --- | --- | --- | --- | --- | --- |
| Trait* | *ABC* | BF100 | LD100 | ADG | FCR | *ABC* | BF100 | LD100 | ADG | FCR |
| ***ABC*** | 0.04<br>(0.01) | 0.19<br>(0.16) | -0.02<br>(0.16) | 0.59<br>(0.17) | 0.30<br>(0.17) | 0.03<br>(0.016) | -0.31<br>(0.21) | -0.24<br>(0.23) | 0.39<br>(0.23) | 0.37<br>(0.20) |
| **BF100** | 0.01<br>(0.02) | 0.57<br>(0.04) | - | - | - | -0.03<br>(0.02) | 0.42<br>(0.04) | - | - | - |
| **LD100** | -0.04<br>(0.02) | - | 0.46<br>(0.04) | - | - | -0.03<br>(0.02) | - | 0.23<br>(0.03) | - | - |
| **ADG** | -0.09<br>(0.02) | - | - | 0.45<br>(0.04) | - | -0.06<br>(0.01) | - | - | 0.33<br>(0.04) | - |
| **FCR** | 0.10<br>(0.02) | - | - | - | 0.32<br>(0.04) | 0.06<br>(0.02) | - | - | - | 0.26<br>(0.04) |

*ABC*: area between the perturbed and unperturbed growth curves; BF100: backfat thickness at 100kg; LD100: *longissimus dorsi* thickness at 100kg; ADG: average daily gain during control; FCR: feed conversion ratio.
\* Standard errors of correlations are given in parentheses.

## Discussion

Although the performance of on-farm fattening pigs has improved over the last decades, phenotypic expression of certain traits remains below their genetic potential. In this context, obtaining reliable estimates of growth potential (unperturbed) and resilience over the fattening period in large populations is a challenge in actual swine breeding conditions. In a perspective of quantifying swine resilience and as an attempt to identify indicator traits for this complex trait, here we described a modelling approach based on pig BW registered routinely by AFS in station conditions: *ad-libitum* feeding, high sanitary level, controlled temperature. Even if conditions are optimal, animals have to face to macro (heat stress, disease outbreak) and micro (social hierarchy, AFS mechanic problem) environmental perturbations that modify expression of growth potential. The modelling approach was tested on different swine breeds and their genetic contribution was analyzed in each one of them. Our modelling approach can further facilitate a real implementation at large scale in pig breeding systems.

### Pretreatment and validation of data registered by AFS

A prerequisite for the linkage of animal data to precision livestock farming systems is through animal identification systems, such as RFID, that are automated and affordable both for the farmer and breeder (22). The development of AFS not only increases the convenience and control of the feeding process, it also allows a precision phenotyping. This development was made possible by the amount of data registered by these devices. These devices routinely record the individual identification, date, age, daily frequency of feeder visits, timing and duration of the visits, FI, and BW (23). However, unlocking the potential of new technology for precision livestock farming requires a deep understanding of how to manage the huge amount of data. Within this framework, data pre-treatment procedures to guarantee high quality data are essential as a first step to exploit the available information. Understanding the data and identifying the main data quality issues require deep data exploration (Figure 1), because modelling approaches are strongly dependent on data quality.

### Quantifying animals’ response to perturbations

Developing models that are able to capture perturbations during the fattening period is a challenge in swine breeding industry. In recent years, the development of more frequent data acquisition and more sophisticated statistical methods have allowed modelling approaches to focus explicitly on perturbations. Revilla *et al*. (11) focused on piglets BW change induced by the weaning event to propose an index to quantify animal robustness during this critical phase. Such a study is based on the modeling of growth, by using with the Gompertz– Makeham function, following an identified disturbance: the weaning. This was shown to correlate with a number of health-related parameters. Nguyen-Ba *et al*. (9) developed a data analysis procedure to detect the impact of perturbations on FI in growing pigs. These two studies aim to analyze and quantify the consequences of an identified disturbance. In the context of our study, pigs can be subjected to different perturbations at different scales depending on the groups: temperature, social hierarchy, health situation. Our model approach does not include an explicit representation of the perturbations and thus differ from other approaches in which the number of perturbations and its duration are either fixed and known (11, 24) or are to be estimated (9, 10). In this study, we described a combined model approach to extract, in a two-step mathematical model approach, perturbed and unperturbed individual growth curves over the pig-fattening period. The Gompertz function (17) was chosen as it is suitable to describe the potential growth of pigs in non-limiting conditions (18, 25). It needs only two parameters, with biological meaning, that can be estimated simply from data (25). The assumption was that the resulting model is an approximation of the theoretical growth rate of the animals not experiencing any perturbation (unperturbed model). The second step characterizes the perturbed growth curve that reflects the production permitted by the farm environment and captures different types of perturbations. With this two-step mathematical model and by comparing the unperturbed and perturbed model a very informative parameter was created, the *ABC* parameter, which gives an estimate of the degree of resilience (11) over the pig-fattening period. Animals can be ranked according to the values of this parameter, with this ranking being an indication on the magnitude of the perturbation and animal resilience. In this case, an *ABC* value parameter closer to zero, means good animal resilience properties. With respect to interpretation, an *ABC* parameter of zero could mean either that the animal is perfectly resilient or that it did not experience any kind of environmental perturbation. In this study, an important hypothesis has been made, we consider that, on average, all animals are subjected to the same perturbations, and so the *ABC* parameter really indicates the resilience response. With this resilience indicator, animals can be ranked based not only on the measured production level, but also on their capacity to cope with perturbations. This kind of approach opens the perspective to use this information for breeding selection. Our hypothesis has however the limitation that we cannot guarantee that all animals are subjected equally to the perturbations. A key challenge is to extend the model to account for the specific perturbations that the individual animals face. Integration of observational data and precision livestock farming technologies are alternatives to explore in future work. For our case study, the interest of genetic analysis is to make it possible to estimate the individual resilience potential by estimating the impact of the environment in which the animal was fattened.

Here Pie and Pie NN lines presented a lower average mean score of parameter *ABC*, −28.89% and −20.48% respectively, compared with Du line (Table 2, Figure 3C). The objective of this comparison is not to conclude that one breed copes better than as other but to illustrate the potential to include a resilience indicator in the selection index. In this scenario, the Du line has a higher level of improvement in terms of selection response to resilience.

### Resilience trait in the breeding objective

The response to societal concerns (*e.g*., antibiotics use, viability, etc) and the need to identify pigs that adapt to diverse and changing environmental conditions make essential that resilience traits, or their indexes, are included in the breeding objective (26). Two pre-requisites to the success of this approach are: a practical and accurate quantitative definition of this resilience trait, and a positive selection response measured with the heritability estimation. The inclusion of heritabilities of functional traits and their feasibility in the breeding objective has been reported (27). In this context, the genetic improvement of resilience traits, maximizing the bottom line instead of performance in a single trait, could be beneficial for the total system profitability (28).

Undoubtedly, directly including resilience traits in future selection criteria will depend on having quantifiable traits that can be recorded cost-effectively and reliably on the large number of animals that are necessary for a breeding program. The estimated heritabilities found in this study are low, ranging from 0.03 to 0.04, suggest that selection for this trait would result in a limited positive selection response. However, the favorable genetic correlations observed between resilience index (*ABC*), and ADG or FCR indicate that gains in both traits can be achieved at the same time, if resilience traits are properly included in the selection criteria. It means that an increase of the resilience index (= a decrease of *ABC*) is globally positively correlated to a genetic improvement of feed efficiency and FCR. Conversely, *ABC* is genetically correlated with growth (ADG), which could be interpreted as that an increase in the genetic potential for ADG increases the risk of a greater deviation of this potential in case of perturbation/stress, that is to say a loss of resilience. Although accuracies of estimates are low, the trends in these correlations must be taken into account in the choice of the weighting applied on each trait of the global index. One difficulty is to define what weighting to give to this resilience index in order to propose a breeding objective balanced with the production traits. Berghof *et al*. (2) proposed a first approach of estimating an economic value of resilience index based on the reduction of time to manage alerts and observations. Beyond the economic value, this approach answers to environmental and societal concerns, that are difficult to quantify.

## Conclusions

This study describes a method to quantify individual resilience during the pig-fattening period, by modelling routine BW measures registered by AFS. In addition, we have identified low to moderate genetic relationship between a resilience indicator and important phenotypic traits in swine production. The heritabilities found for the proposed resilience indicator are low but gives opportunity to be considered as a selection criterion and thus improve resilience. This first approach to building a resilience index, based on an analysis of the growth pattern could be enriched by the inclusion of observations of the environment (health observations, room temperature) and the concomitant analysis of feeding behavior (FI or feeding duration).

## List of abbreviations

ABC: Area between curves
ADG: average daily gain
AFS: automatic feeding system
BF: backfat thickness
BF100: backfat thickness at 100kg
BW: body weight
Du: Duroc
FCR: feed conversion ratio
FI: feed intake
IW: initial weight
LMM: linear mixed model
LD: longissimus dorsi
LD100: longissimus dorsi thickness at 100kg
Pie: Piétrain
Pie NN: Piétrain Français NN Axiom line
PRRS: porcine reproductive and respiratory syndrome
RFID: radio frequency identification
TFI: total feed intake
WG: weight gain
WM: non-null weights
WT: individual testing

## Declarations

### Ethics approval and consent to participate

All animal procedures were performed in accordance with French Animal Welfare legislation. All procedures regarding animal handling and treatment were approved by AXIOM Genetics.

### Consent for publication

Not applicable.

### Availability of data and materials

Data and the Scilab source codes of the procedure described in this article are available under request for academic purposes on the Zenodo data repository (doi: /10.5281/zenodo.4109395).

### Author’s contributions

NCF, RMT, LFG, GL, and MR conceived and designed the study. LFG and GL managed the phenotype recording at the farm. MR and RMT analyzed the data. MR wrote the first draft with input from NCF, RMT and GL. All authors read and approved the final manuscript.

## Acknowledgements

The authors are grateful to AXIOM Genetics for the provision of the data. The authors also thank the personal at AXIOM’s farm at *La Garenne* for their implication for the generation of animals measurements, and their skilful technical assistance. Version 5 of this preprint has been peer-reviewed and recommended by Peer Community In Animal Science (https://doi.org/10.24072/pci.animsci.100008).

## Supplementary material

**Additional File 1 Table S1.**

**Format**: .xlsx

**Title**: Fixed effects in linear mixed models.

**Additional File 2 Figure S1.**

**Format**: .tiff

**Title**: Comparison of body weight density plots before (A) and after (B) applying data cleaning procedure in Pie NN line.

**Description**: In A1 and B1 plots each point represent the median of the individual daily body weight registered by the AFS during the pig fattening period. A2 and B2 are smoothed color density representations of a scatterplot. Shaded areas are constructed to illustrate the density of points falling into each part of the plot allowing for an intuitive visualization of very large datasets. A zoom in the density scatter plot before (A3-B3) and after (A4-B4) 100 days of individual age is illustrated.

**Additional File 3 Figure S2.**

**Format**: .tiff

**Title**: Comparison of body weight density plots before (A) and after (B) applying data cleaning procedure in Du line.

**Description**: In A1 and B1 plots each point represent the median of the individual daily body weight registered by the AFS during the pig fattening period. A2 and B2 are smoothed color density representations of a scatterplot. Shaded areas are constructed to illustrate the density of points falling into each part of the plot allowing for an intuitive visualization of very large datasets. A zoom in the density scatter plot before (A3-B3) and after (A4-B4) 100 days of individual age is illustrated.

