## Supplementary figures and images for "Quantifying growth perturbations over the fattening period in swine via mathematical modelling"

### Additional_File2_Figure_S1

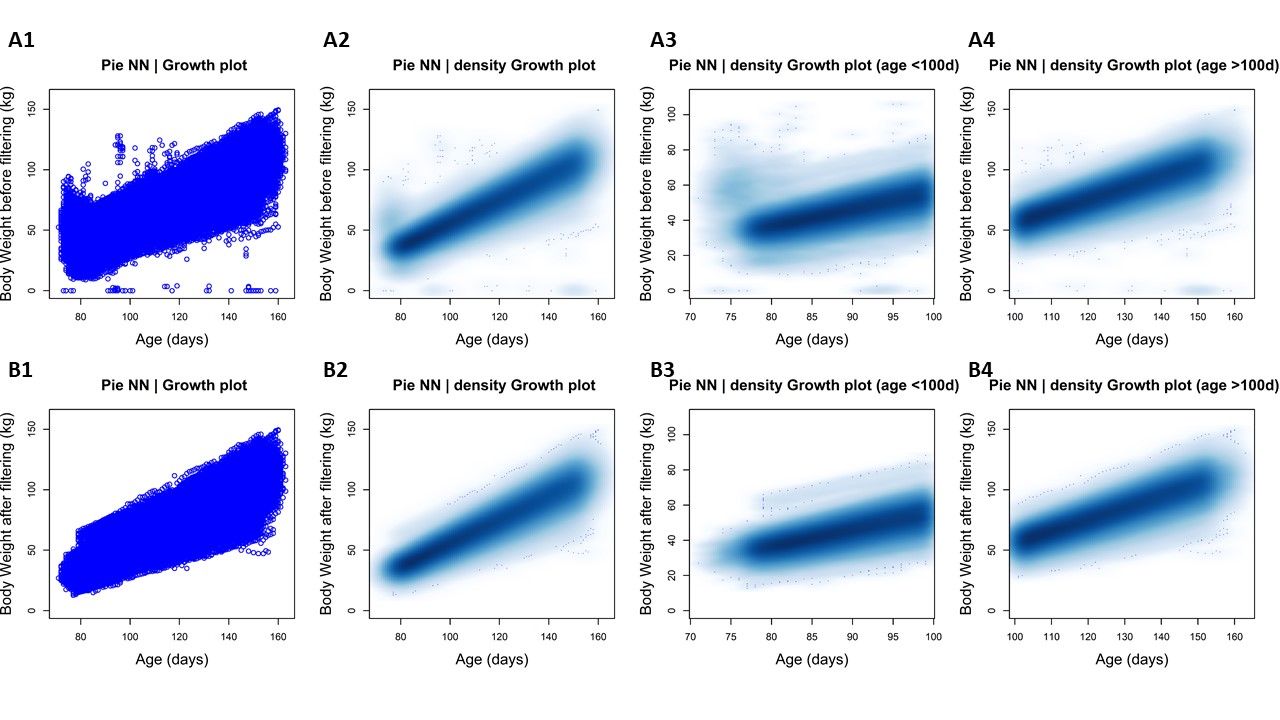

### Additional_File3_Figure_S2

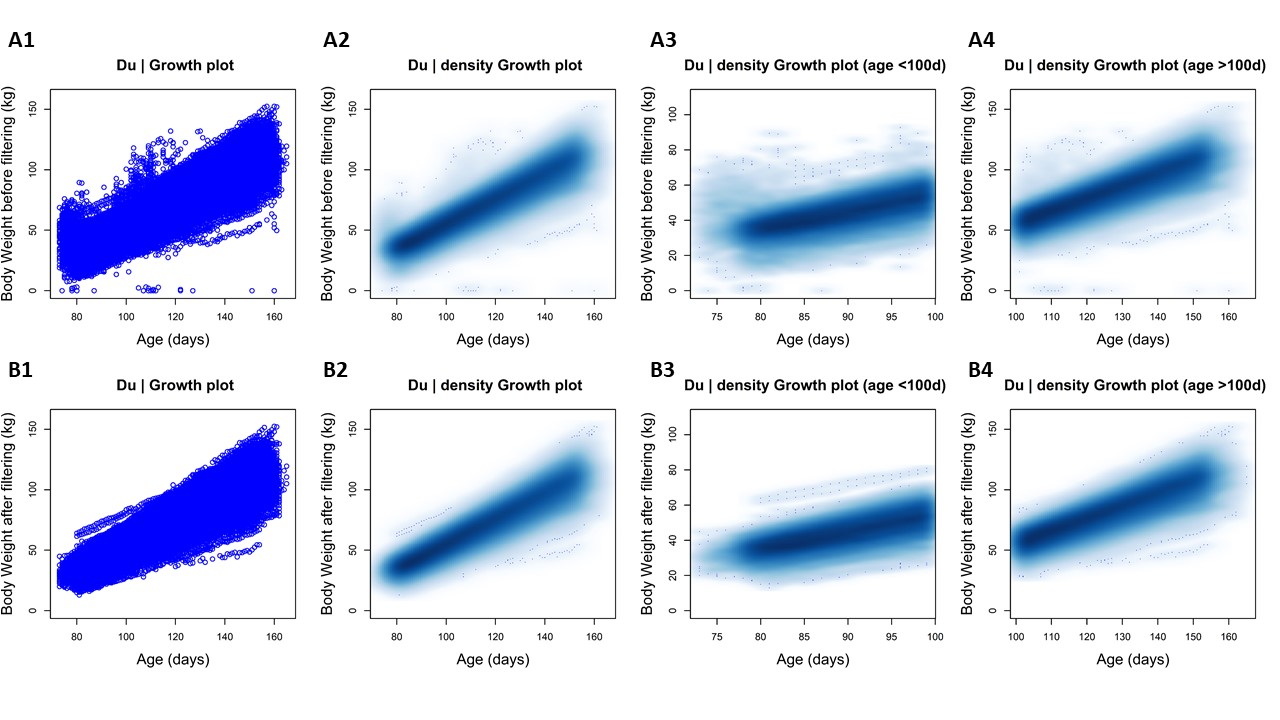
